# Direct detection of mRNA expression in microbial cells by fluorescence *in situ* hybridization using RNase H-assisted rolling circle amplification

**DOI:** 10.1101/729616

**Authors:** Hirokazu Takahashi, Kyohei Horio, Setsu Kato, Toshiro Kobori, Kenshi Watanabe, Tsunehiro Aki, Yutaka Nakashimada, Yoshiko Okamura

## Abstract

Meta-analyses using next generation sequencing is a powerful strategy for studying microbiota; however, it cannot clarify the role of individual microbes within microbiota. To know which cell expresses what gene is important for elucidation of the individual cell’s function in microbiota. In this report, we developed novel fluorescence *in situ* hybridization (FISH) procedure using RNase-H-assisted rolling circle amplification to visualize mRNA of interest in microbial cells without reverse transcription. Our results show that this method is applicable to both gram-negative and gram-positive microbes without any noise from DNA, and it is possible to visualize the target mRNA expression directly at the single-cell level. Therefore, our procedure, when combined with data of meta-analyses, can help to understand the role of individual microbes in the microbiota.

## Introduction

Microbiota consists of several bacterial species, including non-cultivable microbes, via interactions such as nutrients and growth promoting substance. One of the goals of microbiology is to understand the impact of microbiota on the environment, animals including human to marine sponges, and plants including rhizosphere. Thus, meta-analyses such as metagenomics and metatranscriptomics using next generation sequencing (NGS) are powerful strategies for studying microbiota. In particular, high-throughput transcriptome sequencing (RNA-seq) data that contain amount and type of expressed mRNA are an important for understanding the behavior of the microbiota ^1^.

Meanwhile, RNA-seq data cannot clarify the role of individual microbes because complete reference genome of individual microbes is often not available even when metagenomic analysis is performed simultaneously. To clarify the role of individual microbes, single cell genome sequencing of individual microbes in the microbiota has been a trend in recent years ^2, 3^, however high cost restrict many researchers from performing these analysis.. Therefore, fluorescence *in situ* hybridization (FISH) of mRNA is an important method for understanding ‘who is doing what?’ in the microbiota with low cost.

However, compared with the oligo-FISH for 16S rRNA, which is commonly used in microbiology ^4^, mRNA is difficult to visualize by FISH, because the mRNA of interest in a cell has an extremely low copy number compared to the rRNA. In fact, several high-sensitivity FISH methods for mRNA detection have been developed using enzymatic signal amplification such as catalyzed reporter deposition-FISH (CARD–FISH) ^5^ and its derivative methods (e.g. two pass tyramide signal amplification FISH ^6^ and double CARD–FISH ^7^). However, these methods cannot compare mRNA expression level because the signal for these methods diffuses over the entire cell.

Although single-molecule FISH (smFISH) can detect a single mRNA molecule ^8^ even in microbial cells ^9^, this procedure is much expensive than the other FISH method because it needs nearly 50 different detection probes carrying the same fluorophore against one transcript of interest to increase detection sensitivity ^10^. In addition, RNA-seq data without a reference genome, especially from microbiota, are fragmented, and the mRNA sequence of interest is often too short when designing nearly 50 detection probes.

A click-amplifying FISH method, termed ClampFISH, was recently developed to amplify the fluorescence signal ^11^. However, this method is time-consuming and confounded by background noise presumably due to nonspecific hybridization of the probe. Most currently available FISH methods, including RNAscope ^12^ and RollFISH ^13^, rely on simple hybridization of oligonucleotide probes to mRNA sequences. Therefore, these methods cannot clearly distinguish between fluorescent signals originating from mRNA and genomic DNA, especially in microbes. The prokaryotic mRNA is an unspliced RNA; thus, the oligonucleotide probes cannot be set at the exon-exon junction to distinguish between genomic DNA and mRNA, which commonly used in reverse-transcription (RT)-PCR for detection of eukaryotic mRNA.

FISH using rolling circle amplification (RCA), which uses complementary DNA (cDNA) generated from mRNA by RT, has a potential to detect a single mRNA molecule in a eukaryotic cell ^14^. This method makes it possible to distinguish between mRNA and genomic DNA with high probability, because it is difficult to initiate RCA reaction using the genomic DNA as a primer. The RCA product (RCP) contains a hyper tandem-repeat of a DNA sequence complementary to a padlock probe (PLP). The RCP eventually becomes a platform on which a sufficient amount of the detection probe can hybridize. Therefore, single fluorescent probe for detection allows this method to be available at approximately 50-fold lower cost per gene as compared with the conventional smFISH. However, for the detection of microbial mRNA, the exonuclease treatment of cDNA to produce initiation point of RCA reaction will be difficult to optimize for each individual mRNA considering the operon structure, especially using the fragmented sequence data from RNA-seq. For these reasons, a cost-effective and user-friendly FISH method that detects only the target mRNA in a microbial cell is needed.

Recently, we developed RNase H-assisted RCA (RHa-RCA) ^15^. The PLP used in this procedure can be set without full-length sequence information of target RNA; therefore, PLP can be set even with a fragmented mRNA sequence such as RNA-seq data. In addition, this procedure can detect only RNA. Here we demonstrate a novel FISH method based on RHa-RCA for the visualization of mRNA expression level in microbial cells at a single-cell level.

## Results and Discussion

Figure 1 shows the scheme and workflow for our method of visualization of expressing mRNAs of interest in microbial cells. After cell fixation and permeabilization, RHa-RCA–FISH consists of five reaction steps (**Fig. 1A**): (i) PLP hybridization to a sequence of interest in the target mRNA molecule; (ii) circularization of hybridized PLP by SplintR ligase; (iii) producing nick site in the hybridized mRNA by RNase H; (iv) RCA using phi29 DNA polymerase to create RCP from the nick site; and (v) finally, detection probe hybridization to visualize RCP.

**Figure 1.**
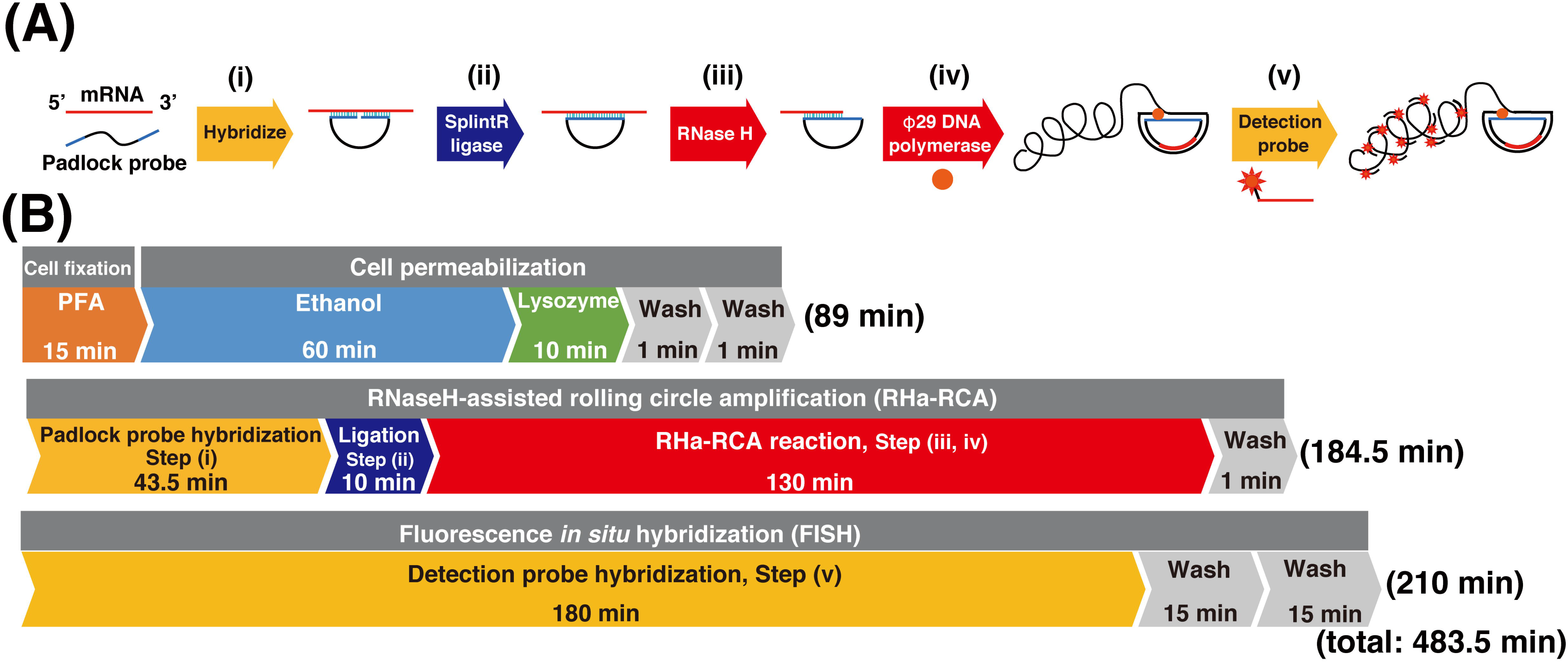
Sheme and workflow for visualization of mRNA in microbial cells. by RNase H-assisted RNA-primed rolling circle amplification (RHa-RCA–FISH). (A) Schematic representation of RHa-RCA–FISH procedure. (B) Workflow of the procedure and required time for *in situ* visualization of mRNA expression.

Using this method, it is possible to complete all steps except fluorescence microscopic observations in approximately 8 h (**Fig. 1B**). Typically, we perform up to step (iv) on the first day and then store the samples at 4°C, as RCPs are stable at this temperature. On the following day, step (v) and microscopy were performed.

At first, we have tested whether RHa-RCA–FISH detects mRNAs of green fluorescent protein (GFP) expressed in *Escherichia coli* cell. *E. coli* BL21(DE3) transformed with a plasmid carrying GFP gene (pET-AcGFP) ^16^ was cultured by inducing GFP expression through the addition of isopropyl-β-d-thiogalactopyranoside (IPTG), whereas the *E. coli* without the addition of IPTG were maintained as negative control (non-induced cell). The non-induced cells were cultured with 2% glucose to inhibit leaky expression of GFP mRNA. The GFP expression was confirmed by fluorescence microscopy using the harvested cells 2 h after the addition of IPTG **(Supplementary Fig. 1)**. Considering that mRNA expression was observed before protein expression, the cells harvested 1 hour after addition of IPTG were used for RHa-RCA–FISH.

No fluorescent signal was observed in non-induced cells, even though the cells were also transformed by pET-AcGFP (**Fig. 2A**). In contrast, fluorescent signals were clearly observed in GFP-induced cells (**Fig. 2B**). This result clearly shows that RHa-RCA–FISH specifically detected GFP mRNA molecules and not the GFP DNA in the vector. In addition, the fluorescence signals formed spot-like shapes (**Fig. 2C**), as observed in microbial cells using smFISH ^9, 10^. Unfortunately, bacterial cells are too small to distinguish each RCP, so it was impossible to determine the exact number of spots in the constrained space of these cells. To count the exact number of spots in a microbial cell, the use of super-resolution microscopy such as Stochastic Optical Reconstruction Microscopy (STORM) is required ^17^.

**Figure 2.**
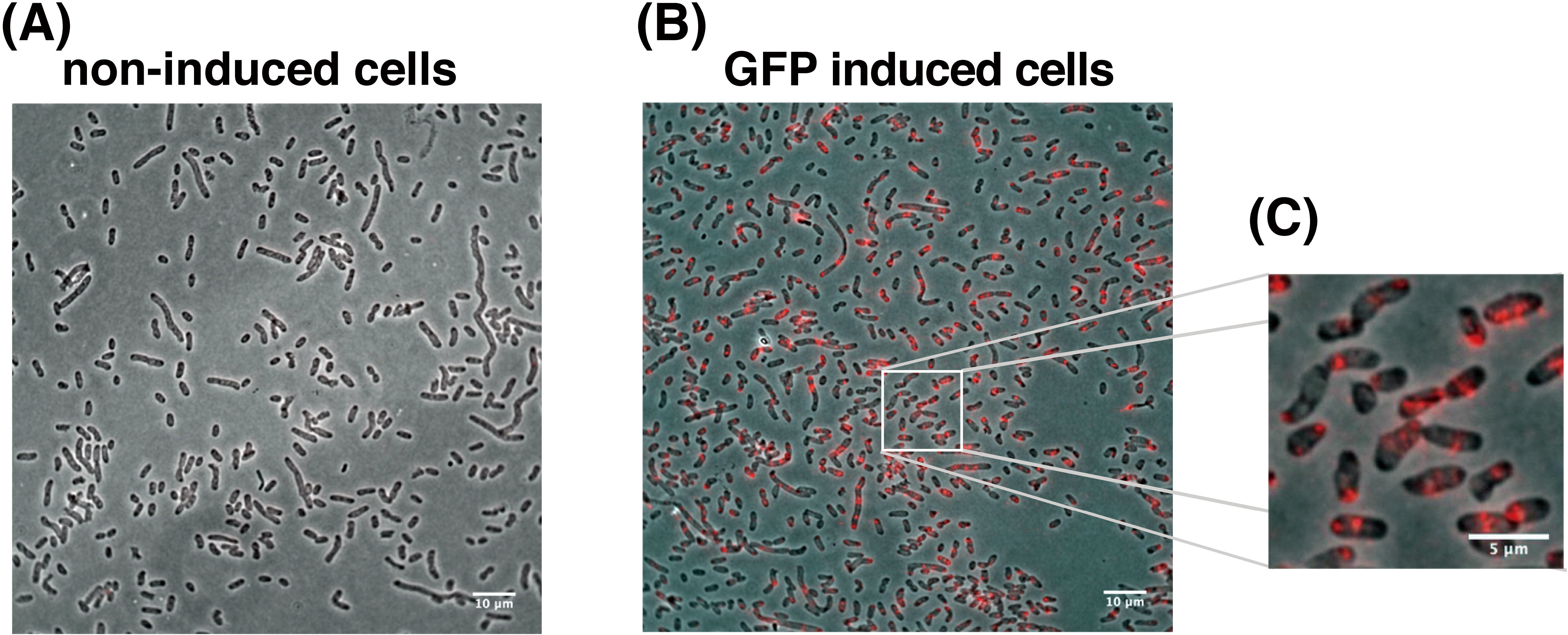
FISH detection of GFP mRNA in *E*. *coli* cells. (A) Detection of GFP mRNA in non-induced *E*. *coli* cells carrying a GFP expression vector and (B) in GFP-induced cells. Scale bar, 10 µm. (C) Magnified image of box in (B). Scale bar, 5 µm. An overlay of the phase contrast (grayscale) and Alexa-568 labeled probes (red) targeting the RCP from GFP mRNA is shown.

Because *E. coli* is a gram-negative bacteria, the utility of RHa-RCA–FISH in microbiota will be limited if this method cannot be performed using the same protocol for gram-positive bacteria, which have a thick peptide glycan coat. Therefore, next we investigated whether RHa-RCA–FISH is applicable to *Brevibacillus choshinensis* as control of gram-positive bacteria.

Using *B. choshinensis* harboring a plasmid carrying the DsRed gene under control of the native P2 promoter, we observed fluorescence from the DsRed protein starting at 12 h after the start of the culture. The fluorescence of DsRed gradually increased, reaching a maximum at 72 h (**Fig. 3, upper images**).

**Figure 3.**
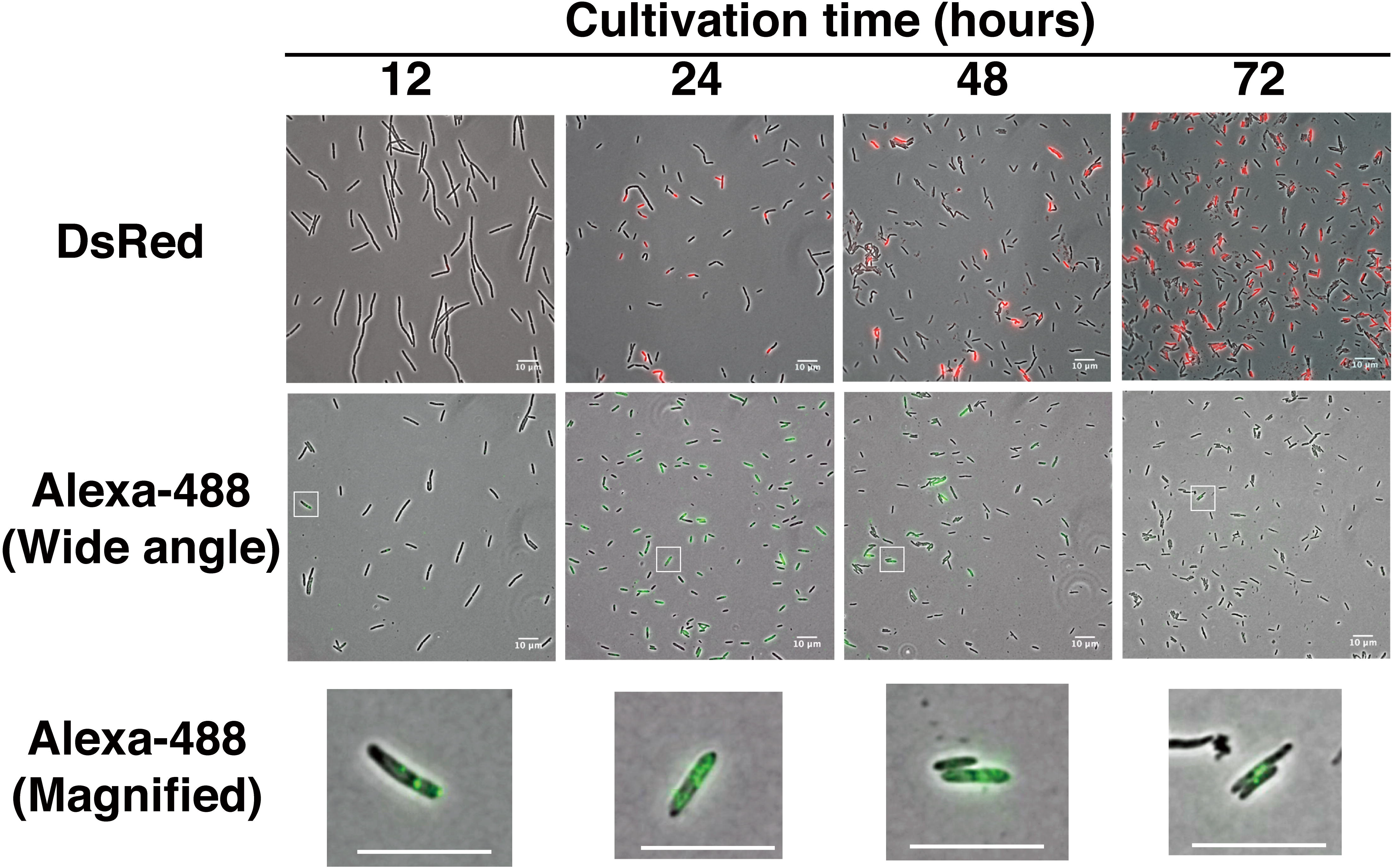
FISH detection of DsRed mRNA in *B*. *choshinensis* as a Gram-positive bacterium. Upper images show fluorescence of DsRed protein expressed in *B*. *choshinensis*; middle images show the signal of detection probes using FISH; lower images show the magnified image of the square box in middle images. The cells in the upper and middle images are not from the same sample because the protein is denatured by the FISH procedure. Overlays of the phase contrast (grayscale), DeRed protein (red), and Alexa-488 labeled probes (green) targeting the RCP from DsRed mRNA are shown. Scale bar, 10 µm.

On the other hand, the fluorescent signal from DsRed mRNA by RHa-RCA–FISH was first observed 12 h after culture initiation, reaching a maximum at 24 h and then decreasing (**Fig. 3, middle images**). These results indicate that RHa-RCA–FISH can be performed following the same protocol even for gram-positive bacteria.

Interestingly, the signal in *B*. *choshinensis* varied greatly between individual cells (**Fig. 3, middle images**), while the fluorescent signals from GFP mRNA in *E. coli* showed minimal variation between cells (**Fig. 2B**). The reason the DsRed mRNA expression level varied between individual cells may be that mRNA expression from the P2 promoter occurs during cell wall synthesis ^18^, which varies with the cell cycle stage. In addition, it was likely that the number of signals could be counted in cells with low mRNA abundance (**Fig. 3, lower images**). These results indicate that RHa-RCA–FISH have a potential to compare the amount of mRNA expression at the single-cell level.

Finally, we performed our protocol for simultaneous detection of both GFP and DsRed mRNA in suspension containing *E. coli* and *B*. *choshinensis* cells. As a result, red fluorescence (Alexa -568) and green fluorescence (Alexa -488) were observed from separate cells (**Fig. 4**). Although both the strains have similar cell shapes that cannot be discriminated under the microscopy we were able to distinguish between the two strains based on the species-specific fluorescence from labelled probes. This result indicates that RHa-RCA–FISH can simultaneously detect multiple specific mRNAs, even in microbiota containing both gram-positive and gram-negative bacteria.

**Figure 4.**
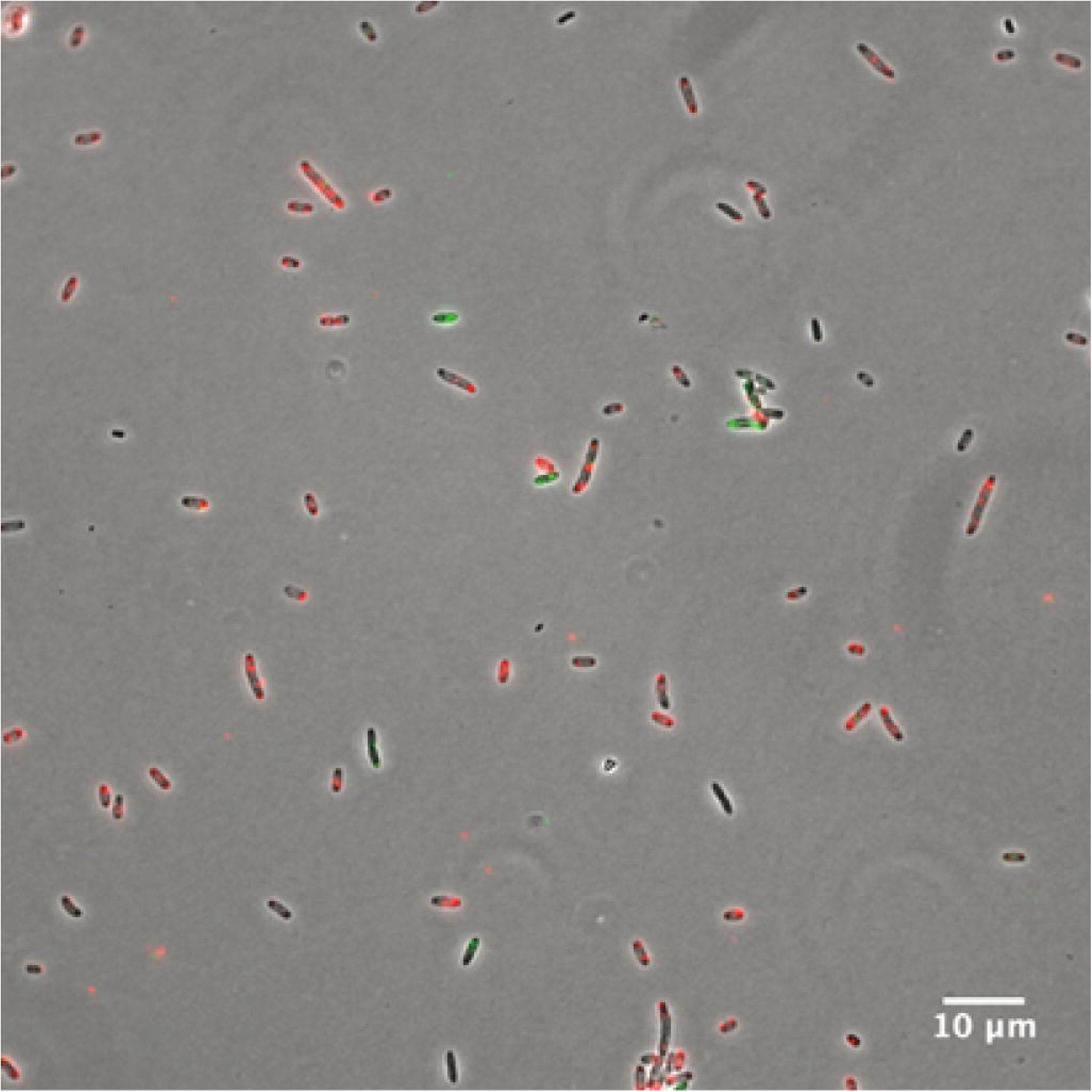
Simultaneous detection of GFP and DsRed mRNA in a mixture of *E. coli* and *B*. *choshinensis* cells. An overlay of the phase contrast (grayscale) and Alexa-568 labeled probes (red) targeting the RCP from GFP mRNA and Alexa-488 labeled probes (green) targeting the RCP from DsRed mRNA are shown. Scale bar, 10 µm.

Here we report that our method for *in situ* analysis of mRNA expression level in both gram-positive and gram-negative bacteria has potential to detect differences in amount of the mRNA between individual microbial cells without background noise from genomic or plasmid DNA. In this report, we performed RHa-RCA–FISH using both gram-positive and gram-negative bacteria in liquid culture; thus, the samples used for RHa-RCA–FISH contain almost no impurities other than bacterial cells. In contrast, the samples that many researchers want to study, such as feces, soil, and symbiont, contain impurities other than bacterial cells. However, we believe that RHa-RCA–FISH can be perform for samples containing impurities because oligo-FISH for 16S rRNA has been performed on many samples including feces ^19^, soil ^20^ and symbiont ^21^. However, it is important to examine the method of fixing and permeabilization according to the state of the sample to be observed.

In this report, we have demonstrated that RHa-RCA–FISH will be able to compare the amount of mRNA expression in rod-shaped bacterial cells. However, whether the expression levels of mRNA can be compared greatly depends on the physical size and shape of the microbial cells. It is difficult to compare the expression level in the cells of small cocci with a commonly used fluorescence microscope. The FISH signal and the cells were almost the same size, therefore, the mRNA expression level was not revealed in *Streptococcus thermophilus*, which used for fermentation of yogurt (data not shown). Thus, we believe that super-resolution microscopes such as STORM ^17^ will become essential in microbiology.

We used RNase H to digest RNA/DNA-hybrid but not DNA/DNA-hybrid molecules, therefore, our method can clearly distinguish between mRNA and DNA even prokaryotic mRNA remains unspliced. In addition, because the PLP used in this method can be set anywhere in the mRNA sequence ^15^, the PLP can be set even in the RNA-seq data which may contain short sequence. Thus, we believe that our method for visualizing RNA molecules directly within cells could help understand the role of individual microbes in the microbiota when combined with the data of meta-analyses.

## Material and methods

### Padlock probe and detection probe

Each PLP position is shown in **Supplementary Figs. 3–4**. The PLP and detection probe sequences are shown in **Supplementary Table 1**. All PLP and PCR primers were purchased from Eurofins genomics (Ebersberg, Germany). Alexa labeled detection probes were purchased from Japan Bio Services Co., LTD.(Asaka, Saitama, Japan).

### Expression of GFP mRNA in *E*. *coli*

The *E*. *coli* strain BL21 (DE3) (Novagen, Merck Millipore, Darmstadt, Germany) carrying pET-AcGFP was grown in LB medium (1% tryptone, 0.5% yeast extract, 1.0% NaCl, and 50 µg/mL ampicillin) for 14–16 h at 30°C with shaking. The cultures were diluted 1:1000 in fresh LB medium and incubated at 30°C with shaking. Growth was monitored by the measurement of the optical density at 660 nm (OD_660_). When the OD_660_ was approximately 0.6, GFP expression was induced by the addition of IPTG (final concentration, 0.5 mM). The GFP-induced cells for RHa-RCA–FISH were harvested 1 h after IPTG addition. GFP-induced and non-induced cells were collected in 1.5 mL tubes by centrifugation at 10,000 × *g* for 2 min at 4°C. Cells were immediately suspended in saturated ammonium sulfate (SAS) solution to inhibit RNase activity and store at 4°C before use. This treatment would keep the total RNA in the cells stable without degradation for at least 2 weeks (**Supplementary Fig. 2)**.

### Construction of expression vector for *B*. *choshinensis* and *in vitro* transcription

The coding sequence of DsRed was isolated from the pDsRed-monomer vector (Clontech/TaKaRa Bio, Ohtsu, Shiga, Japan) by digestion with the appropriate restriction enzymes. The resulting fragment was cell-free cloned into pNI-His (Takara Bio) for expression in *B*. *choshinensis* cells as pNI-DsRed or pET-21d (Novagen) for *in vitro* transcription as pET-DsRed. Detailed procedures for cell-free cloning are described in the **Supplementary Material and Methods**.

### Expression of DsRed in *B*. *choshinensis*

The *B*. *choshinensis* strain HPD31-SP3 (Takara Bio) carrying pNI-DsRed was grown in 2SYF medium (20.0 g/L fructose, 40.0 g/L Phytone Peptone (Becton, Dickinson, and Co., Franklin Lakes, NJ), 5.0 g/L Ehrlich bonito extract (Kyokuto Pharmaceutical Co.LTD, Tokyo, Japan), and 0.15 g/L CaCl_2_·2H_2_O) containing 50 µg/mL neomycin for 12, 24, 48, or 72 h at 30°C with shaking. DsRed protein expression was confirmed by fluorescence microscopy. The cells were harvested in 1.5 mL microcentrifuge tubes by centrifugation at 10,000 × *g* for 2 min at 4°C. Cells were suspended in SAS and stored at 4°C before use.

### Cell fixation and permeabilization

The stored cells in SAS solution (600 µL) were transferred to a 1.5 mL tube and centrifuged at 20,000 × *g* for 1 min at 4°C to remove the SAS solution. The cell pellet was resuspended in 300 µL of 4% paraformaldehyde and incubated at room temperature for 15 min to fix the cells. After centrifugation, the cell pellet was resuspended with 300 µL of 70% ethanol and incubated at room temperature for 60 min to dehydrate the cells. After centrifugation under the same conditions to remove 70% EtOH, the cell pellet was resuspended with 262.5 µL of 1× TE buffer. Then, 37.5 µL of lysozyme solution (200 µg/mL in 50% glycerol) was added to the cell suspension solution and incubated at room temperature for 10 min to digest the cell wall. After centrifugation at same conditions to remove the lysozyme solution, the cells were washed twice with 300 µL of 1× phosphate-buffered saline (PBS). After centrifugation, the cells were resuspended with 14 µL of µDW and transferred to 0.2 mL of PCR tube to prepare the RCA reaction.

### FISH using RHa-RCA

The cells were mixed with 20 pmol of PLP in a buffer containing 20 mM Tris-acetate (pH 7.5), 10 mM magnesium acetate (MgAc), and 50 mM potassium glutamate (KGlu) in a final volume of 20 µL. The PLP was hybridized by incubation at 95°C for 1 min, slowly cooling to 30°C over 30 min, and incubation for 10 min at 30°C. To the cells was added 10 µL of ligation mixture (20 mM Tris-acetate [pH 7.5], 10 mM MgAc, 1.2 mM ATP, 50 mM KGlu, 10 mM dithiothreitol, and 25 units of SplintR ligase (New England BioLabs, Ipswich, MA)), followed by incubation at 37°C for 10 min to seal the PLP. In our previous report ^15^, the ligase was inactivated by heating, but this step was omitted in this study to shorten the reaction time and simplify the procedure. The detection sensitivity was not affected (data not shown). The RHa-RCA reaction was started by mixing 30 µL of the ligated mixture with 20 µL of a reaction mixture containing 20 mM Tris-acetate (pH 7.5), 10 mM MgAc, 80 mM ammonium sulfate, 10 mM KGlu, 2.0 mM deoxynucleoside triphosphate, 0.004 units of pyrophosphatase (New England BioLabs), 0.06 units of RNase H (BioAcademia, Osaka, Japan), and 500 ng of DNA-free phi29 DNA polymerase (Kanto Chemical, Tokyo, Japan). The mixture was incubated at 30°C for 2 h followed by enzyme inactivation at 65°C for 10 min.

The RCA reaction mixture containing the cells was transferred to a 1.5 mL tube. After centrifugation at 20,000 × *g* for 1 min at 4°C, the cells were washed with 1× PBS at room temperature, suspended in 2× saline sodium citrate, and transferred to a 1.5 mL black tube. After addition of the Alexa-labeled oligonucleotide, the cell suspension was incubated at 37°C for 3 h to hybridize the oligonucleotide to RCP. After centrifugation at 20,000 × *g* for 1 min at 4°C, the cells were washed twice with 1× PBS at 37°C for 15 min. The cells were suspended in 1× PBS and stored at 4°C in 1.5 mL black tubes before observation by microscopy.

RCA and hybridization were performed in bench-top cleanroom ^22^ to prevent contamination that could cause nonspecific hybridization..

### Imaging and analysis

A drop of cell suspension hybridized with the detection probe was placed on 1% agarose pads containing 1× PBS. Images of each sample were taken on a fluorescence microscope (Nikon ECLIPSE E600 for *E*. *coli* and Nikon ECLIPSE Ti2-E for *B*. *choshinensis*, Tokyo, Japan) equipped with a phase-contrast objective CFI PlanApo DM 100×(Nikon) and an ORCA-Flash4.0 V3 camera (Hamamatsu Co., Shizuoka, Japan). Typically, the RHa-RCA–FISH, GFP, and DsRed fluorescence images were taken using an exposure time of 100 msec. The images were analyzed using ImageJ software (NIH).

## Acknowledgments

We thank the members of the Okamura lab for helpful discussions. This study was supported in part by the Step-Up Support Program for KAKENHI (Grant-in-Aid for Scientific Research) of Hiroshima University and Adaptable and Seamless Technology transfer program through target-driven R&D (A-STEP) of JST [No. AS2311331E].

## Contributions

H. T, T. K, Y. N, and Y. O conceived of this study and designed the experiments. K. H performed nearly all of the experiments. K. H and S. K collected and analyzed the FISH images. T. K and Y. O obtained the necessary financial support. H. T, K. H, and Y. O prepared the initial draft of the manuscript. T. K, S. K, T. A, K. W, and Y. N revised the manuscript. All authors contributed to and have approved the final manuscript.

