## Supplementary file for "Direct detection of mRNA expression in microbial cells by fluorescence *in situ* hybridization using RNase H-assisted rolling circle amplification"

Supplementary Figure, Table, and Material and Methods

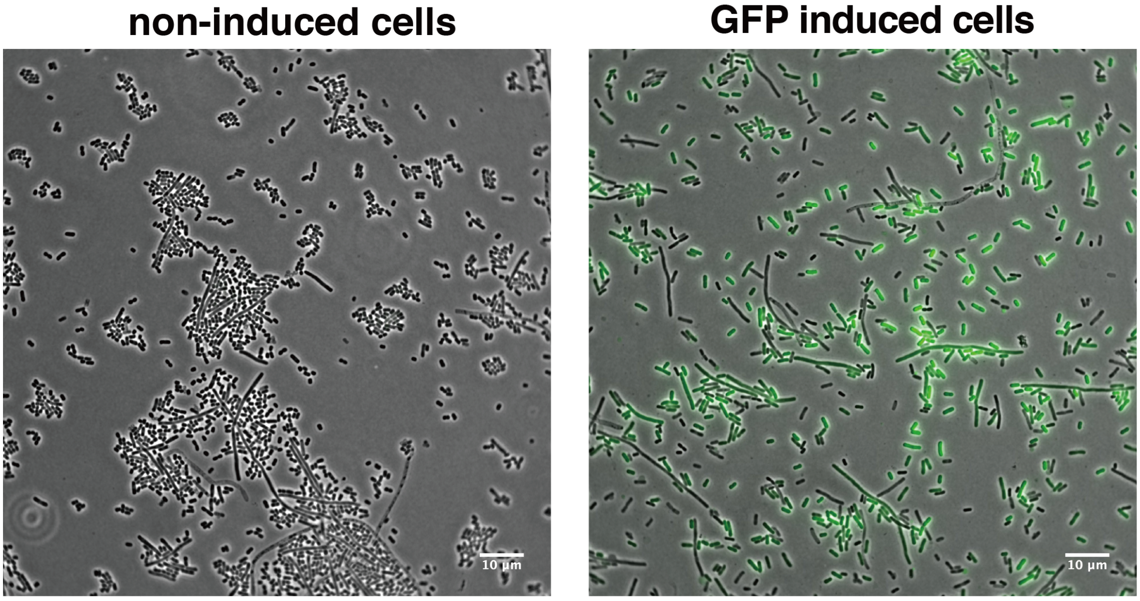


**Supplementary Figure 1**

Observation of GFP fluorescence in *E.coli* cells. The GFP induced cells were harvested at 2 h after addition of IPTG. The non-induced cells were harvested without addition of IPTG. The non-induced cells were cultured with 2% glucose to inhibit leaky expression of GFP mRNA. Scale bar, 10-µm.

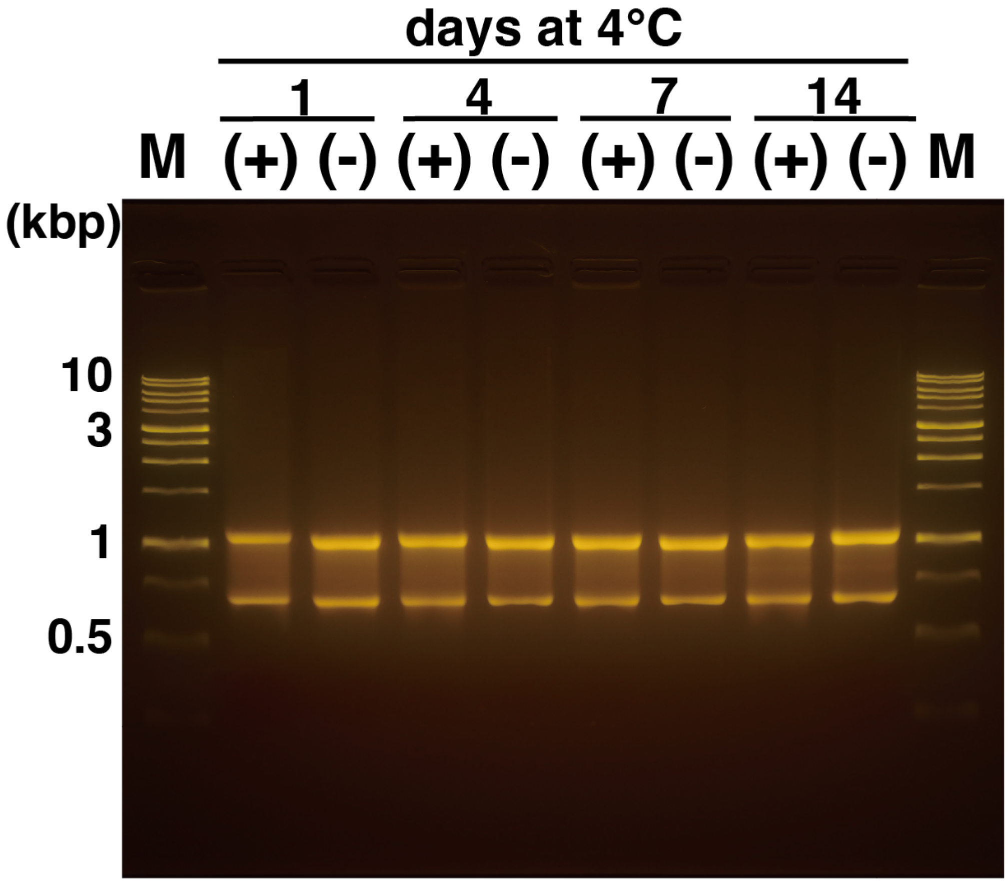


**Supplementary Figure 2**

Agarose gel analysis of extracted total RNA from *E.coli* cells suspended in saturated ammonium sulfate solution and stored at 4°C. After denatured with formamide (final conc. 47.5%), the RNA samples were loaded into well of 1% TAE agarose gel. The numbers above the lanes indicate the number of days stored at 4 ° C. (+), GFP induced cells; (-), non-induced cells; M, 1 kbp DNA ladder. RNA and DNA were visualized by staining using UltraPower$^{TM}$ DNA/RNA Safe Dye and a blue-light transilluminator. Since the TAE gel is not a denatured gel for RNA, the length of the double-stranded DNA size marker is not informative.

```
  1 ATGGTGAGCAAGGGCGCCGAGCTGTTCACCGGCATCGTGCCCATCCTGATCGAGCTGAAT
 61 GGCGATGTGAATGGCCACAAGTTCAGCGTGAGCGGCGAGGGCGAGGGCGATGCCACCTAC
121 GGCAAGCTGACCCTGAAGTTCATCTGCACCACCGGCAAGCTGCCTGTGCCCTGGCCCACC
181 CTGGTGACCACCCTGAGCTACGGCGTGCAGTGCTTCTCACGCTACCCCGATCACATGAAG
241 CAGCACGACTTCTTCAAGAGCGCCATGCCTGAGGGCTACATCCAGGAGCGCACCATCTTC
                          ***************###############
301 TTCGAGGATGACGGCAACTACAAGTCGCGCGCCGAGGTGAAGTTCGAGGGCGATACCCTG
361 GTGAATCGCATCGAGCTGACCGGCACCGATTTCAAGGAGGATGGCAACATCCTGGGCAAT
421 AAGATGGAGTACAACTACAACGCCCACAATGTGTACATCATGACCGACAAGGCCAAGAAT
481 GGCATCAAGGTGAACTTCAAGATCCGCCACAACATCGAGGATGGCAGCGTGCAGCTGGCC
541 GACCACTACCAGCAGAATACCCCCATCGGCGATGGCCCTGTGCTGCTGCCCGATAACCAC
601 TACCTGTCCACCCAGAGCGCCCTGTCCAAGGACCCCAACGAGAAGCGCGATCACATGATC
661 TACTTCGGCTTCGTGACCGCCGCCGCCATCACCCACGGCATGGATGAGCTGTACAAGTGA
```

#### Supplementary Figure 3

Padlock probe position for GFP mRNA (red color words). The sequence of GFP shows from ATG (Met) to TGA (stop codon). “*” are indicate a 5’arm region, and “#” are indicate a 3’-arm region of padlock probe for GFP (P3).

```
  1 ATGACCATGATTACGCCAAGCTTGCATGCCTGCAGGTCGACTCTAGAGGATCCCCGGGTA

 61 CCGGTCGCCACCATGGACAACACCGAGGACGTCATCAAGGAGTTCATGCAGTTCAAGGTG

121 CGCATGGAGGGCTCCGTGAACGGCCACTACTTCGAGATCGAGGGCGAGGGCGAGGGCAAG

181 CCCTACGAGGGCACCCAGACCGCCAAGCTGCAGGTGACCAAGGGCGGCCCCCTGCCCTTC

241 GCCTGGGACATCCTGTCCCCCCAGTTCCAGTACGGCTCCAAGGCCTACGTGAAGCACCCC

301 GCCGACATCCCCGACTACATGAAGCTGTCCTTCCCCGAGGGCTTCACCTGGGAGCGCTCC

361 ATGAACTTCGAGGACGGCGGCGTGGTGGAGGTGCAGCAGGACTCCTCCCTGCAGGACGGC

421 ACCTTCATCTACAAGGTGAAGTTCAAGGGCGTGAACTTCCCCGCCGACGGCCCCGTAATG

481 CAGAAGAAGACTGCCGGCTGGGAGCCCTCCACCGAGAAGCTGTACCCCCAGGACGGCGTG

541 CTGAAGGGCGAGATCTCCCACGCCCTGAAGCTGAAGGACGGCGGCCACTACACCTGCGAC
                                                 ***************

601 TTCAAGACCGTGTACAAGGCCAAGAAGCCCGTGCAGCTGCCCGGCAACCACTACGTGGAC
    ###############

661 TCCAAGCTGGACATCACCAACCACAACGAGGACTACACCGTGGTGGAGCAGTACGAGCAC

721 GCCGAGGCCCGCCACTCCGGCTCCCAGTAG
```

#### Supplementary Figure 4

Padlock probe position for DsRed mRNA (red color words). The sequence of DsRed shows from ATG (Met) to TAG (stop codon). “*” are indicate a 5’arm region, and “#” are indicate a 3’-arm region of padlock probe for DsRed (614R).

**Supplementary Table 1**. Sequence of padlock probe and detection probe for GFP and DsRed mRNA.

| Name | Sequence | Length |
|---|---|---|
| GFP padlock probe_P3 | 5'-*AGCCCTCAGGCATGG*ttccttttacgaCCTCAATGCTGCTGCTGTACTACtcttc*TGCGCTCCTGGATGT*-3' | 70 mer |
| Alexa568-detection probe | 5’-Alexa568-CCTCAATGCTGCTGCTGTACTAC-3’ | 23 mer |
| DsRed padlock probe_614R | 5'-*AGTCGCAGGTGTAGTG*tttcttttactcCCTCAATGCACATGTTTGGCTCCtcttt*GTACACGGTCTTGA*-3' | 70 mer |
| Alexa488-detection probe | 5'-Alexa488-CCTCAATGCACATGTTTGGCTCC-3' | 23 mer |

Italic uppercase letters indicate homologous sequence to GFP or DsRed mRNA .Uppercase letters indicate detection probe sequence. The sequence of detection probes were referred to the sequences of Larsson et al.[1, 2] GFP padlock probe_P3 is same sequence in our previously report.[3]

### Supplementary Material and Methods

#### Solutions, mixtures, plastic ware, and DNA

To avoid contamination from laboratory environment, prepared solutions supplied by the manufacturers were used as much as possible in this study. UltraPURE™ distilled water (μDW) were purchased from Thermo Scientific. All homemade solutions and buffers were filtrated and sterilized using a 0.1-μm-pore-size polyethersulfone membrane bottle top filter unit (Nalgene/Thermo Scientific). Similarly, disposable sterile plastic ware was used whenever possible to reduce the likelihood of DNA and RNase contamination. Aerosol-resistant filter tips were purchased from Molecular BioProducts/Thermo Scientific and DNA low binding microcentrifuge tubes and PCR-grade 0.2-ml tubes were obtained from Eppendorf. Black microcentrifuge tubes for shielding from light were obtained from Watson Bio Lab. All solutions and mixtures were prepared using dedicated sets of pipettes in a classified as ISO-1 bench-top cleanroom (KOACH 500F, Koken Ltd)[1] after cleaning using RNase *AWAY* (Molecular BioProducts).

#### Oligonucleotides

All padlock probes were purchased from Eurofins Genomics, Inc. Alexa labelled oligonucleotides were purchased from Japan Bio Services Co., LTD. Each padlock probe position is shown in **Supplementary Fig. 3-4** and the sequences are shown in **Supplementary Table 1**. Thiophosphated random RNA hexamer (6R5S, 5'-rN*rN*rN*rN*rN*rN; *, thiophosphate-linkage; rN, random RNA base)[2] was also purchased from Tsukuba oligo service Co., LTD.

All oligomers were dissolved in 0.1×Tris-EDTA (TE) buffer (1 mM Tris-HCl pH 7.5, 0.01 mM EDTA) in the bench-top cleanroom as described above.

**Construction of expression vector for *B. choshinensis* and *in vitro* transcription**

Basically, all cloning procedure were performed with cell-free cloning method[2] using phi29 DNA polymerase. At first, pDsRed-monomer vector (Clontech) was amplified by multiply-primed rolling circle amplification (MPRCA)[3] using RNA primer (6R5S)[2] to obtain several micrograms of the DNA. The amplified pDsRed was digested by *Nco*I (Takara Bio) with Shrimp Alkaline Phosphatase (SAP, Takara Bio) at 37°C for 1 h following 65°C for 10min to inactivate enzymes. Then, the linearized pDsRed was digested by *Not*I (New England BioLabs) for pET-21d (Novagen) or *Eco*RI (Takara Bio) for pNI-His (Takara Bio) to separate the DsRed fragment (approximately 0.7 kb). Then, each DsRed fragments were isolated by agarose gel electrophoresis and purified with Wizard® SV Gel and PCR Clean-Up System (Promega).

The pET-21d and pNI-His vector were also amplified by MPRCA using 6R5S. The amplified pET-21d was digested by *Not*I (Takara Bio) with SAP at 37°C for 1 h following 65°C for 10min to inactivate enzymes. The amplified pNI-His was digested by *Eco*RI with SAP at 37°C for 1 h following 65°C for 10min to inactivate enzymes. Then, each linearized vector was digested by *Nco*I, and then, isolated by agarose gel electrophoresis and purified with Wizard® SV Gel and PCR Clean-Up System.

The each DsRed fragments and vectors were ligated using T4 DNA ligase (Takara Bio) in 1× CutSmart buffer (New England BioLabs) with 1 mM ATP at 16°C for 30 min following 65°C for 10 min to inactivate enzymes. To remove the linear DNA, each ligated product were treated with Plasmid Safe ATP-Dependent DNase (Epicentre) and Exonuclease I (New England BioLabs) in 1× CutSmart buffer with 1 mM ATP (final conc.) at 37°C for 4 h following 80°C for 20 min to inactivate enzymes. Then, remained circular ligated products were amplified with 6R5S and DNA-free phi29 DNA polymerase[4] (Kanto Chemical) at 37°C for 16 h following 65°C for 10 min to inactivate enzymes according to our previously report[2]. The success or failure of cloning was confirmed by restriction enzyme digestion (single- and double-digestion) and agarose gel electrophoresis.

##### *In vitro* transcription of DsRed mRNA

*In vitro* transcription of DsRed mRNA was performed using a ScriptMAX® Thermo T7 Transcription Kit (Toyobo) and linearized pET-DsRed by *Not*I digestion according to the manufacturer's protocol. After RNase-free DNase I (New England BioLabs) treatment, the transcribed mRNA was then purified using a NucleoSpin® RNA Clean-up XS kit (Macherey-Nagel) according to the manufacturer's protocol. RNA concentrations were determined using a Qubit® fluorometer (Invitrogen/Thermo Fischer Scientific) with Qubit® RNA BR Assay Kit (Invitrogen). The transcribed mRNA was stored at -80°C until required.

##### Confirmation of padlock probe for DsRed by real-time RHa-RCA

The padlock probe (PLP) was checked by real-time RHa-RCA using *in vitro*-transcribed DsRed mRNA according to our previously report[5]. The *in vitro*-transcribed DsRed mRNA was mixed with 250 fmol of a PLP in a buffer containing 20 mM Tris-acetate (pH 7.5), 10 mM magnesium acetate (MgAc), 1 mM ATP, and 50 mM potassium glutamate (KGlu) in a final volume of 9 μl. Hybridization of the padlock probe was facilitated by incubation at 95°C for 1 min followed by immediate cooling to 40°C and incubation for 3 min at 40°C and 10 min at 30°C. After hybridization, 1 μl of SplintR ligase (25 units) was added to the reaction mixture, which was incubated at 37°C for 10 min to seal the PLP. Finally, the RHa-RCA reaction was started by mixing 10 μl of the ligated mixture with 10 μl of a reaction mixture containing 20 mM Tris-acetate (pH 7.5), 10 mM MgAc, 80 mM ammonium sulfate, 10 mM KGlu, 2.0 mM deoxynucleoside triphosphate, 10 mM dithiothreitol, 0.002 units of pyrophosphatase (New England BioLabs), 0.06 units of RNase H (BioAcademia), 2× concentration of SYBR Green II (Invitrogen), and 100 ng of DNA-free phi29 DNA polymerase (Kanto Chemical)[4].

For real-time detection, RHa-RCA reaction was performed in a 96-well PCR plate, and fluorescence signals were measured every 10 min for 2 h with the FAM filter (excitation wavelength: 482 nm, fluorescence wavelength: 536 nm) of the Thermal Cycler Dice Real-Time System II (TP900, Takara Bio). All real-time RHa-RCA were performed in triplicate to estimate the experimental variance.

#### Transformation of *B. choshinensis*

Transformation of *B. choshinensis* was performed according to the manufacturer's protocol. The *B. choshinensis* strain HPD31-SP3 (Takara Bio) was transformed with re-ligated *Eco*RI digested cell-free cloned pNI-DsRed and then grown on MT agar plate (10.0 g/L glucose, 10.0 g/L BBL™ Phytone™ Peptone (Becton, Dickinson and Co.) 5.0 g/L 35% Ehrlich Bonito Extract (Kyokuto Pharmaceutical Industrial Co.), 2.0 g/L yeast extract (Nakarai Tesque) , 10 mg/L $FeSO_4 \cdot 7H_2O$, 10 mg/L $MnSO_4 \cdot 4H_2O$, 1 mg/L $ZnSO_4 \cdot 7H_2O$, 4.1 mg/L $MgCl_2 \cdot 6H_2O$ and 1.5% agar) containing 50 μg/mL neomycin to select clones transformed with pNI-DsRed.
